# Cell coupling compensates for changes in single-cell Her6 dynamics and provides phenotypic robustness

**DOI:** 10.1101/2022.12.02.518899

**Authors:** Parnian Doostdar, Joshua Hawley, Elli Marinopoulou, Robert Lea, Veronica Biga, Nancy Papalopulu, Ximena Soto Rodriguez

## Abstract

*her6* is a zebrafish ortholog of *Hes1*, known for its role in maintaining neural progenitors during neural development. Here, we characterise the population-level effect of altering Her6 protein expression dynamics at the single-cell level in the embryonic zebrafish telencephalon. Using an endogenous *Her6:Venus* reporter and 4D single-cell tracking, we show that Her6 oscillates in neural telencephalic progenitors and that fusion of a protein destabilisation domain (PEST) to Her6:Venus alters its expression dynamics causing most cells to downregulate Her6 prematurely. However, in PEST mutants, a higher proportion of cells exhibit Her6 oscillations and while expression is reduced in most cells, some cells express Her6 at wild-type levels resulting in increased heterogeneity of Her6 expression in the population. Despite the profound differences in the single-cell Her6 dynamics, differentiation markers do not exhibit major differences early on, while an increase in differentiation is observed at later developmental stages (*vglut2a, gad1* and *gad2*). At the same time, at late stage the overall size of the telencephalon remains the same. Computational modelling that simulates changes in Her6 protein stability reveals that the increase in population Her6 expression heterogeneity is an emergent property of finely tuned Notch signalling coupling between single cells. Our study suggests that such cell coupling provides a compensation strategy whereby a normal phenotype is maintained while single-cell dynamics are abnormal, although the limit of this compensation is reached at late developmental stages. We conclude that in the neural progenitor population, cell coupling controls Her6 expression heterogeneity and in doing so, it provides phenotypic robustness when individual cells lose Her6 expression prematurely.

## Introduction

Oscillations of transcription factor (TF) proteins are powerful transmitters of cellular information that direct cellular outcomes in various biological contexts including DNA damage response, cell proliferation and development (Isomura and Kageyama, 2014). Oscillatory expression of basic helix-loop-helix (bHLH) Hairy and Enhancer of split (HES in mammals and Her in zebrafish) transcriptional inhibitors are critical in driving the vertebrate segmentation clock (Aulehla et al., 2008) (Rhode et al., 2021). With advances in molecular engineering and generation of luminescent/fluorescent promoter reporters and later endogenous knock-ins, HES/Her proteins have also been found to be oscillatory in the developing central nervous system (CNS) where they also drive oscillations of proneural factors such as DELTA, NGN2 and ASCL1 (Imayoshi et al., 2013a). These oscillations, with periodicities within the range of hours (i.e. ultradian), have been shown to be critical for correct progression of CNS development.

The functional relevance of TF oscillations has been examined with a wide range of experimental approaches to alter their dynamic expression. These include methods like exogenous TF expression from a ubiquitous promoter (Marinopoulou et al., 2021; Sabherwal et al., 2021) or even direct optogenetic manipulation of dynamics (Imayoshi et al., 2013b; Isomura et al., 2017; Shimojo et al., 2016). Many of the manipulations of oscillatory dynamics aimed to uncover function, have been based on the molecular requirements for generating oscillatory expression (Novak and Tyson, 2008). Specifically, oscillations are often based on a negative feedback loop coupled with biological time delays, associated with transcription and translation, as well as instability of mRNA and protein. As such, in CNS development, changes in mRNA stability/translation have been achieved by mutating or blocking microRNA binding sites in the endogenous gene (Bonev et al., 2012) (Soto et al., 2020) while changes in time delays have been achieved by changing the length of introns (Ochi et al., 2020; Shimojo et al., 2016). Collectively, these functional studies have shown that HES/Her oscillations are important in dictating cell-fate decisions and transitions. These include maintaining mouse telencephalic neural progenitors in development (Imayoshi et al., 2013a; Shimojo et al., 2008), transition from progenitor to differentiation states in zebrafish hindbrain (Soto et al., 2020) as well as neural stem-cell exit from quiescence (Marinopoulou et al., 2021; Sueda and Kageyama, 2020).

Another intriguing layer of complexity in HES/Her dynamics is their upstream regulation by Notch signalling, enabling cell states and decisions to be integrated between neighbouring cells. However, the tissue level organisation of oscillators is far from intuitive and requires specific investigation, combining theory and real-time observation, in order to be understood. Using such methods, it has been recently reported that in the developing mouse CNS, HES5 oscillators organise in microclusters of synchronised cells which, in addition, are spatially periodic and temporally dynamic (Biga et al., 2021). This complex organisation requires weak cell-cell coupling via Notch signalling and in turn controls the spatiotemporal dynamics of neuronal differentiation (Biga et al., 2021; Hawley et al., 2022). However, whether such tissue level organisation has additional roles during perturbed development is not currently known.

In order to characterise single-cell and tissue level characteristics of the HES/Her oscillators in perturbed tissue, we have focused on *her6*, the *Hes1* orthologue in the optically superior vertebrate model, zebrafish. The aim of our work presented in this paper is twofold within the context of the developing zebrafish telencephalon. First, we wanted to investigate the changes in Her6 single-cell oscillatory dynamics by alterations of a less explored molecular requirement for oscillations, protein stability. Second, we sought to examine the tissue-level response to the altered single-cell dynamics and to do so in relation to the observed phenotype. To reduce protein stability, we have taken advantage of the CRISPR/Cas9 technology to insert a Venus fluorescent protein fused to a protein destabilisation domain (PEST) domain at the C-terminus of the endogenous Her6. PEST sequences are widely accepted signals for making proteins susceptible to rapid proteolysis and can exert this effect when fused to other proteins (Aulehla et al., 2008; Li et al., 1998; Ninov et al., 2012; Rogers et al., 1986).

We report that Her6 oscillates in some zebrafish telencephalic progenitors and that, surprisingly, destabilising Her6 results in an increase in the proportion of progenitors that show oscillatory Her6 expression. As expected, destabilising Her6 also results in a pronounced and premature downregulation of Her6 during development and overall, a lower level of Her6 protein in the majority of these telencephalic progenitors. However, counterintuitively, interspersed with these, some cells retained high levels of Her6 expression. Together, the opposing effects in Her6 levels, lead to increased expression heterogeneity in the population. This is also reflected in increased cell-cell variability between neighbouring cells in the presence of destabilised Her6.

Computational modelling suggests that this increased variability of Her6 expression in neighbouring cells and interspersion of high-low cells, is a tissue-level property that emerges out of the coupling of Her6 dynamics between cells. Surprisingly, despite the altered Her6 expression at single-cell level, the telencephalon was phenotypically normal early on (24hpf) in terms of size and differentiation, with some increased neurogenesis observed at late stages. We conclude that when Her6 is destabilised, a tissue-level communication serves to partially “rescue” Her6 levels in some cells and to increase the proportion of cells with oscillatory Her6 expression, thus providing some phenotypic robustness to the population in the face of altered molecular dynamics in single cells.

## Results

### *Her6:Venus* is a faithful reporter of proliferative embryonic neural progenitors in zebrafish telencephalon

Her6 has been well-characterised in the development of the hindbrain and hypothalamus (Pasini et al., 2001; Scholpp et al., 2009) but its expression pattern and dynamics in the developing telencephalon have not been extensively explored. To characterise Her6 expression dynamics in zebrafish telencephalon we used the *Her6:Venus* transgenic line (*HV)*, that generates the endogenous protein fusion Her6:Venus (HV)) described by (Soto et al., 2020) (**Fig. 1A**). Soto et al. have demonstrated that fusion of Venus to Her6 does not affect normal development and faithfully recapitulates the expression of endogenous Her6 in the hindbrain. Here we focused on characterising Her6 in the forebrain using embryos between 17 to 24 hours post fertilisation (hpf) to cover early neurogenesis (**Fig. 1B-D** and **Fig S1**). First, we compared HV protein from homozygote *HV* knock-in embryos with *her6* mRNA from wild-type embryos in the forebrain region and observed a broadly similar HV pattern of expression (**Fig. 1B**). We also conducted double whole mount fluorescent in situ hybridisation (WM FISH) against endogenous *her6* and *venus* in homozygous *HV* knock-in embryos (**Fig. 1C**) which showed broad co-localisation. These findings showed that Her6 is expressed in the rostral and ventral telencephalon and confirmed that HV expression is co-localised with *her6* with no aberrant expression, demonstrating the suitability of the *HV* model for studying Her6 telencephalic expression.

**Figure 1:**
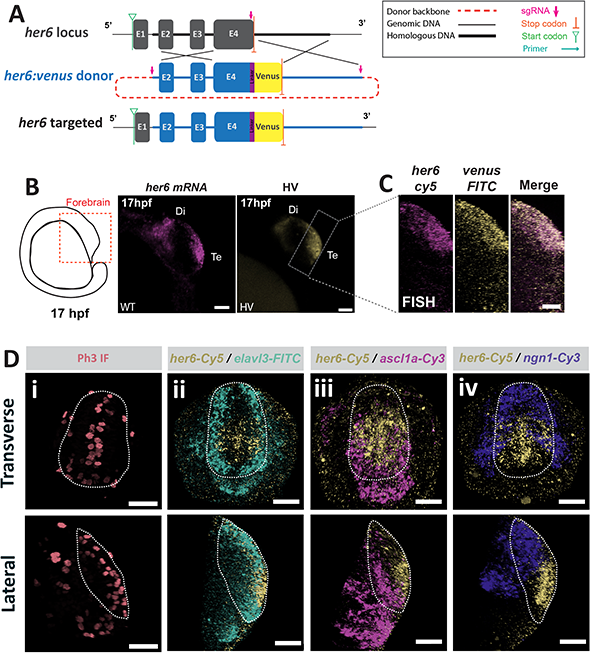
*her6* is expressed in a subpopulation of neural progenitor cells in *Her6:Venus* zebrafish telencephalon. **(A)** Schematic diagram of the *Her6:Venus* knock-in design from (Soto et al., 2020). CRISPR/Cas9-mediated homology directed repair (HDR) (demonstrated by crossing lines) resulted in in-frame incorporation of *venus* at the 3’ end of the endogenous *her6* locus. **(B)** Diagram depicting the zebrafish embryo forebrain at 17hpf (left panel). Maximum intensity projection image of WM FISH of endogenous *her6* in wild type (WT) embryo (mid panel) and live snapshot of Her6:Venus (HV) expression in single Z-plane from an embryo homozygous (HOM) for the *Her6:Venus* knock-in (*HV)* (right panel). Lateral view showing the telencephalon (Te) and diencephalon (Di) (scale bars S0μm). **(C)** Maximum intensity projection images from double WM FISH against *her6* and *venus* mRNAs in 17hpf HOM *HV* embryo. Lateral view of the telencephalon corresponding roughly to the marked region in the right panel in (B) (scale bars 20μm). **(D)** Images of the telencephalon from 24hpf HOM *HV* embryos in transverse (top) and lateral (bottom) views with manually annotated presumptive telencephalon boundaries (dashed contour). **(i)** WM immunofluorescent (IF) staining for Phospho-histone H3 (Ph3) shows mitotic activity at the telencephalic midline. **(ii)** WM FISH against *her6/elalv3* shows *her6* expression media-ventrally and lack of overlap with the bilateral *elav/3* expression. **(iii)** WM FISH against *her6/asc/1* shows expression of *asc/1* posterior to *her6* with no major overlap. **(iv)** WM FISH against *her6/ngn1* shows expression of *ngn1* dorsal to *her6* with no major overlap. (Scale bars 40μm).

Second, we investigated the spatial localisation of *her6* in relation to known markers of proliferation and differentiation **(Fig. 1D)**. *her6* was expressed in proliferating cells, as indicated by antibody staining against the mitotic marker Ph3 expressed at the telencephalic midline, and its expression did not overlap with the early post-mitotic neural marker *elavl3* (Kim et al., 1996) nor the proneural factors *ascl1a* and *ngn1* (Schmidt et al., 2013), suggesting that *her6* expressing cells are indeed proliferative embryonic neural progenitors **(Fig. 1Di-iv)**.

### Her6 telencephalic expression exhibits heterogeneity, due to ultradian oscillations and asynchronous downregulation in single cells

*her6* mRNA can be detected in the presumptive forebrain as early as 11hpf and persists in the developing telencephalon up to at least 30hpf **(Fig. S1)**. To characterise Her6 dynamics in the telencephalon, we performed live imaging between 20-30hpf to capture the time window with the highest rate of neural differentiation while Her6 was still expressed **(Fig. 2A)**. Although the length of the imaging varied, the 20-26hpf interval was always shared between experiments **(Fig. 2B-C)**.

**Figure 2:**
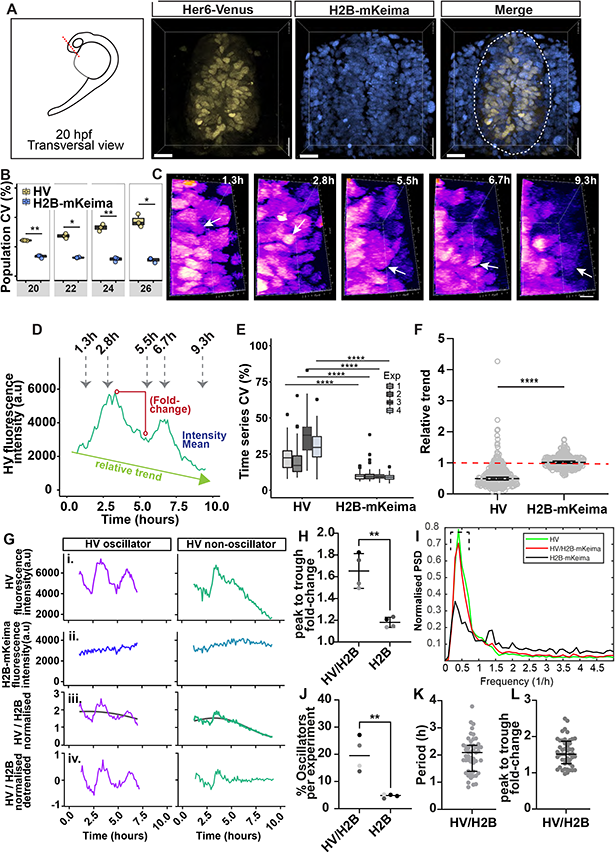
Her6 expression is heterogeneous across the telencephalon in *HV* embryos. **(A)** Schematic diagram showing the transversal view of the telencephalon at 20hpf. Red dotted line indicates the position of live-imaging (left panel). Transversal view of 3D reconstructed confocal live images from 20hpf telencephalon of *HV* embryo showing HV (yellow) and H2B-mKeima (blue) (marking single nuclei). Manual annotation indicates the telencephalon boundaries (dashed contour) (right panel) (scale bars 20μm). **(B)** Coefficient of variation (CV-see Materials and Methods) at population level for HV and H2B-mKeima in *HV* embryos within 20-26hpf. Boxplots indicate the interquartile range and vertical lines the median. Each dot represents the collective CV of all cells for each developmental stage per embryo with an average of 397 cells per developmental stage per embryo. HV and H2B-mKeima values are collected from the same cell (2way-ANOVA with Sidak multiple comparisons; the statistics inside the panels show multiple comparisons). **(C)** 3D reconstructed confocal images depicting a single cell expressing HV (white arrows) over time. Snapshots show HV expression from 1.3 to 9.3 hours following the onset of imaging at 20hpf. Pseudo-colour indicates Venus intensity ranging from blue (low) to white (high). **(D)** HV expression over time of the single cell marked in (C). The arrows indicate the timepoints where the snapshots shown in (C) were acquired. The graph also depicts parameters analysed in other panels. These include: Intensity mean of a single cell over time (blue dashed line), relative trend (calculated by dividing the last timepoint in each time-series to the first) and fold-change (peak-to-trough difference in intensity (red circles)). On X axis 0=20hpf. **(E)** Boxplots comparing the CV for each cell trace over time (denoting relative variability around the intensity mean over time per cell trace) between HV and H2B-mKeima time-series from the same cell. (Boxes indicate interquartile range from 4 biological repeats, vertical lines correspond to median, an average of 76 cells per biological repeat is shown, dots indicate outliers, two-tailed Wilcoxon matched-pairs non-parametric signed rank test). **(F)** Comparison of relative trend between HV and H2B-mKeima obtained from the same cell trace. Relative trend is calculated by dividing the last timepoint in each time-series to the first. Values> 1 indicate upregulation, values = 1 indicate steady expression and values < 1 indicate downregulation. (Boxes indicate interquartile range from 4 biological repeats, vertical lines correspond to median, each dot represents the relative trend of a single time-series (n=260), Mann-Whitney). **(G) (i. left and right, respectively)** Representative examples of oscillatory and non-oscillatory HV time-series with corresponding H2B-mKeima traces **(ii. left and right)** obtained from the same nuclei. **(iii. left and right)** Same HV trace after being divided by H2B-mKeima (HV/H2B normalised). **(iv. Left and right)** Normalised traces after removing the long-term trend (HV/H2B normalised-detrended). **(H)** Comparison of peak-to-trough fold-change in HV/H2B (Her6:Venus/H2B-mKeima) versus H2B (H2B-mKeima) collected in the same nuclei (bars indicate median and interquartile range from 4 biological repeats, dots indicate median per condition from 40 to 80 cells per embryo, statistical test is one-tailed paired t-test). **(I)** Aggregate power spectral density (PSD) obtained from time-series HV, HV/H2B and nuclear control H2B-mKeima collected in the same cells. **(J)** Percentage of oscillators in detrended HV/H2B versus H2B control (bars indicate median from 4 biological repeats from 40 to 80 cells per embryo, each dot indicates median per experiment, statistical test is one-tailed paired t-test). **(K)** Measure of HV period from HV/H2B detrended single cells (bars indicate median and interquartile range from 4 biological repeats of 10-12 cells per embryo, dots indicate individual cells. **(L)** Peak to-trough fold-change in HV oscillators measured from HV/H2B (bars indicate median and interquartile range from 4 biological repeats of 12-12 cells per embryo; dots indicate individual cells). Statistical significance(*): P ≤ 0.05, (**): P ≤ 0.01, (***): P ≤ 0.001, (****): P ≤ 0.0001.

*HV* knock-in embryos were injected with *h2b-mkeima* mRNA for the purposes of segmentation and single-cell analysis (Materials and Methods). Snapshot images from transverse views of the telencephalon at 20hpf showed a heterogenous HV expression among single cells **(Fig. 2A)**. This was confirmed by estimating the Coefficient of Variation (CV), the ratio of standard deviation of nuclear intensities at a specific time to the mean, which was used to measure snapshot heterogeneity of expression in the cell population **(Fig. 2B**, Materials and Methods). HV expression appeared progressively more heterogeneous between 20-26hpf whereas, as expected, this effect was not observed for the stable nuclear marker H2B-mKeima collected from the same nuclei **(Fig. 2B)**. Overall, HV was consistently and significantly more variable than H2B-mKeima at all stages **(Fig. 2B)**.

This increase in heterogeneity during the course of development could be the result of changes in single-cell HV expression over time. For instance, (1) Her6 expression may be fluctuating between low and high values, and/or (2) it may downregulate as development progresses (long-term trend). HV expression was tracked at single-cell level between 20 to 30hpf to generate single-cell fluorescence intensity traces **(Fig. 2C,D)** where we observed fluctuations in protein expression (see example in **Fig. 2C** with matching timestamps in **Fig. 2D)**. To confirm the presence of heterogeneity in HV expression in single-cell traces, we measured the CV in single-cell time-series (Materials and Methods) **(Fig. 2E)**. HV expression had significantly higher variability compared to the nuclear control H2B-mKeima, showing that the population heterogeneity **(Fig. 2B)** arises from changes in HV expression over time at single-cell level **(Fig. 2E)**.

To investigate whether Her6 expression is downregulated over the course of development, we quantified the relative trend in each time-series by dividing the last intensity value to the first (relative trend ratio) **(Fig. 2F)**. H2B-mKeima time-series had relative trend ratios predominantly close to 1 showing steady expression over time whereas the large majority of HV time-series showed downregulation **(Fig. 2F)**. Downregulation was asynchronous with some cells reaching trend ratios far below 1 while others retained levels and remained closer to 1 **(Fig. 2F)**. Overall, the presence of downregulation contributed to the observed Her6 single-cell heterogeneity **(Fig. 2E)**. However, Her6 also exhibited fluctuations in expression overlaid onto a slow-varying downward trend **(Fig. 2D)** so we investigated if these short-term dynamics also contribute to Her6 heterogeneity.

We have previously shown, in hindbrain, that Her6 protein expression at single-cell level can be periodic (oscillatory) or aperiodic (noisy), fluctuating every few hours representing ultradian dynamics (Soto et al., 2020). Thus, we investigated ultradian activity in time-series of HV in single neural progenitors from the telencephalon **(Fig. 2G-L)**. Single cells were tracked to monitor HV **(Fig. 2G i)** and H2B-mKeima **(Fig. 2G ii)** fluorescence intensity over time and HV was normalised to mKeima fluorescence intensity to remove any potential imaging artefacts **(Fig. 2G iii**, see HV/H2B-Materials and Methods). Then, in order to focus on the ultradian dynamics we removed the long-term trends (such as downregulation) in the normalised traces **(Fig. 2G iv**, see HV/H2B normalised-detrended). In the zebrafish telencephalon, both oscillatory and non-oscillatory HV expression was observed **(Fig. 2G**, and additional examples **Fig. S2A)**. We used the Hilbert transform to identify peaks and troughs in the Venus signal and measure peak-to-trough fold-change as a mean to describe amplitude of intensity fluctuations **(Fig. S2B)** (Materials and methods) (Manning et al., 2019). The Her6 amplitude in all cells ranged between 1.5x to 1.8x and was more prominent than noisy fluctuations observed in nuclear H2B-mKeima **(Fig. 2H, fold-change)**. Frequency analysis indicated that the Venus signal has a prominent peak around 2.5h, indicative of presence of oscillatory activity **(Fig. 2I)**. The dominant frequency peak was closely reproduced by the HV/H2B signal and as expected, appeared severely dampened in the non-oscillatory H2B-mKeima observed in the same cells **(Fig. 2K)**. Thus, fluctuations in HV are not due to imaging artifacts such as nuclear movement.

To distinguish periodic (oscillatory) expression from stochastic aperiodic (non-oscillatory) expression, single cell HV/H2B traces were analysed using a statistical method of stochastic oscillatory activity detection using Gaussian processes (Materials and Methods; (Phillips et al., 2017; Soto et al., 2020). HV oscillatory cells represented 15 to 30 % of the total neural progenitor cells analysed **(Fig. 2J)**. Due to an imposed False Discovery Rate, only a small number of H2B-mKeima traces were found oscillatory (approximately 5%, **Fig 2J)** and appeared strikingly different from the HV oscillators **(Fig. 2G, black trace** and **Fig S2A)**. Importantly, the percentage of oscillators detected in HV/H2B was significantly higher than the observed in nuclear H2B-mKeima in the same cells **(Fig. 2J)**. The detected oscillators showed a median of 1.9h period ranging between 1 to 3h in good agreement with previous reports (Soto et al., 2020). The amplitude of oscillations ranged between 1.5x to 2.5x **(Fig. 2L, fold-change)**, similar to overall amplitude in the population **(Fig. 2H, fold-change)**. This indicated that both oscillatory and non-oscillatory ultradian dynamics contribute to gene expression heterogeneity.

### Decreasing HV protein stability increases the number and amplitude of oscillators and promotes downregulation

We hypothesised that changes in Her6 protein properties, such as its rate of degradation, would affect its dynamic behaviour with a concomitant effect on neural differentiation. To test this hypothesis, we aimed to alter Her6 protein properties by creating an in-frame fusion of the endogenous Her6 protein with Venus fluorescent protein plus a protein destabilising PEST domain at the C-terminus, generating the *Her6:Venus:PEST (HVP)* zebrafish line **(Fig. 3A;** Materials and Methods and **Fig. S3A,B)**. The spatial localisation of the *her6* domain in *HVP* was comparable to *HV*, in the rostral tip of the developing telencephalon **(Fig. S3C, D)**. We also validated this construct by conducting WM FISH against endogenous *her6* and *venus* in homozygous *HVP* knock-in embryos **(Fig. S3E, F)**. To confirm that the insertion of the PEST domain destabilises the HV protein, we carried out protein half-life experiments **(Fig. S4)**. As previously shown, the Her6 protein half-life is shorter (11min, (Soto et al., 2020)) relative to its mouse orthologue HES1 (22.3 minutes) (Kobayashi et al., 2015). We expected that by adding one PEST domain, this would reduce even further the Her6 protein half-life (potentially in the range of few minutes) and would therefore make it technically more difficult to estimate in zebrafish. As an alternative, we used human mammalian cells to express and compare the half-lives of the HV and HVP proteins, where we were expected this to be in the range of hours according to what has been reported before (Sabherwal et al., 2021) **(Fig. S4A)**. MCF7 cells expressing HV or HVP were treated with cycloheximide, and the protein degradation rate was estimated in both conditions (Materials and Methods). Expression of the HVP protein declined more rapidly than HV **(Fig. S4B-C)** and the HVP protein half-life was significantly reduced when compared to HV **(Fig. S4D)**

**Figure 3:**
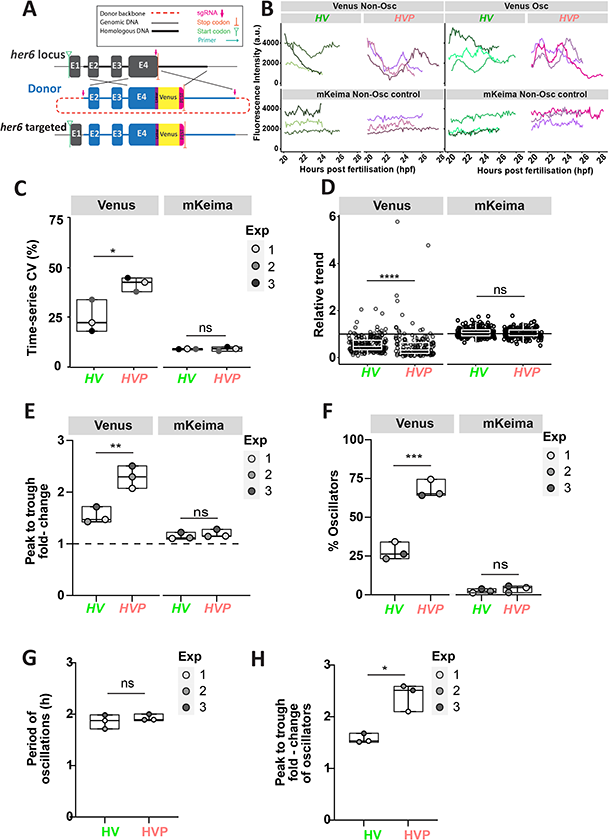
Her6 expression dynamics in single cells are altered when the protein is destabilised. **(A)** Schematic diagram of the *Her6:Venus-PEST (HVP)* knock-in design. CRISPR/Cas9-mediated HOR resulted in the insertion of the Kl cassette including Venus and PEST into the endogenous locus, a more detailed description is presented in **Fig. S3A,B**. **(B)** Representative examples of Venus oscillatory (Venus Osc) and Venus non-oscillatory (Venus Non-Osc) and their equivalent mKeima time-series obtained from *HV* and *HVP* embryos between 20-30hpf (traces shown from 3 biological repeats). **(C)** Comparison of time-series CV (denoting relative variability around the intensity mean over time) from raw single-cell Venus and H2B-mKeima time-series observed in the same cells (boxes indicate median and interquartile range from 3 biological repeats, dots indicate median per embryo, statistical test is two-tailed t-test, ns: P=0.9381). **(D)** Comparison of relative trend of Venus and H2B-mKeima raw time-series between *HV* and *HVP* embryos (relative trend is calculated by dividing the last timepoint in each time-series to the first, values > 1 refers to upregulation, values = 1 refers to steady expression and values < 1 refers to downregulation, boxes indicate median and interquartile range from 3 biological repeats, dots represent individual cells expressing Venus or H2B-mKeima (*HV* n=199, *HVP* n=165), statistical test is Mann Whitney, ns: P=0.4207). **(E)** Comparison of the mean fold-change in oscillators expressing raw Venus and their respective mKeima between *HV* and *HVP* embryos (boxes indicate median and interquartile range from 3 biological repeats, dot represent median in *HV*: 40-80 cells and *HVP*: 40-70 cells per experiment, statistical test is two-tailed t test, ns: P=0.4798). **(F)** Comparison of the percentage of oscillators between *HV* and *HVP* embryos. Cells expressing Venus (HV and HVP respectively) and mKeima (H2B-mKeima from the respective *HV* and *HVP* embryos) (boxes indicate median and interquartile range from 3 biological repeats, dots represent median of *HV*: 40-80 cells and *HVP*: 40-70 cells per experiment, statistical test is paired two-tailed t test, ns: P=0.3609). **(G)** Period of oscillatory HV versus HVP expression (boxes indicate median with interquartile range from 3 biological repeats, dots represent median of HV: 10-20 cells and HVP: 30-34 cells per embryo, statistical test is paired two-tailed t-test, ns: P=0.3165). **(H)** Peak-to-trough fold-change in amplitude of oscillatory HV versus HVP expression (boxes indicate median with interquartile range from 3 biological repeats, dots represent median of HV: 10-20 cells and HVP: 30-34 cells per embryo, statistical test is paired two-tailed t-test). Statistical significance (ns): P >0.05, (*): P≤0.05, (***): P≤0.001, (****): P≤0.0001.

To test the effect of changing the Her6 protein stability, we compared HV with HVP protein dynamics in single cells between 20-28hpf. Statistical detection using Gaussian Processes (Materials and methods) revealed that similar to HV dynamic protein expression, both oscillating and non-oscillating Her6:Venus expression are present in *HVP* **(Fig. 3B)**. To examine Her6 fluctuations more quantitatively, we analysed the time-series CV, which revealed that HVP time-series were significantly more variable across all experiments when compared to HV **(Fig. 3C** and **Fig. S4E)**. The increased heterogeneity in *HVP* was in part due to differences in the long-term trend, quantified as relative trend ratio (final intensity divided by initial intensity), where HVP showed more drastic downregulation relative to HV **(Fig. 3D** and **Fig. S4F)**. At the same time, few observations of HV and HVP showed upregulation (trend ratio>1) with remarkable examples in HVP reaching values above maximum HV **(Fig. 3D** and **Fig. S4F)**.

Overall, we found that the peak-to-trough fold-change values in HVP time-series were consistently higher than HV across experiments **(Fig. 3E)**. We also observed a significant increase in the percentage of oscillators in HVP when compared to HV, **(Fig. 3F;** approximately 70% and 25%, in HVP versus HV, respectively). The oscillatory period remained unaltered **(Fig. 3G** and **Fig. S4G)** while the amplitude significantly increased from approximately 1.6x in HV to 2.5x in HVP **(Fig. 3H** and **Fig. S4H)**. As expected, the nuclear aperiodic control H2B-mKeima collected in the same cells showed no differences between HV and HVP conditions **(Fig. 3C-F** and **Fig S4E,F)** and an acceptable FDR of 5% **(Fig. 3F)**.

Overall, these quantitative comparisons showed that the increased single-cell variability in HVP compared to HV is a composite of increased Her6 downregulation, an unexpected rise in the proportion of cells with oscillatory expression and an increase in peak-to-trough fold-changes that denotes increased amplitude.

### Decreasing Her6 stability increases spatial heterogeneity in single-cell protein expression in the telencephalon

Following the characterisation of Her6 protein dynamics at single-cell level, we set out to investigate protein expression differences at tissue level. Snapshots from live imaging demonstrate that fewer cells express Her6 in *HVP* and their expression appears more variable between neighbouring cells than in *HV* at 20, 24 and 26hpf **(Fig. 4A)**. To quantify this difference, we used the H2B-mKeima to detect nuclei in the imaged domain of the telencephalon, in both *HV* and *HVP*, regardless of their Venus expression. To separate Venus(+) and Venus(−) cells a detection threshold was set for each embryo based on background measurements (see Materials and Methods). The analysis reflected lower abundance of cells expressing Venus for the duration of live imaging in *HVP*, quantified as the proportion of Venus(+) cells over the total number of cells expressing H2B-mKeima **(Fig. 4B** and **Fig. S5A)**. We also observed a tendency for faster decrease of Venus(+) cells in *HVP* when compared to *HV* **(Fig. S5B)** however, this was not reproducible across all experiments.

**Figure 4:**
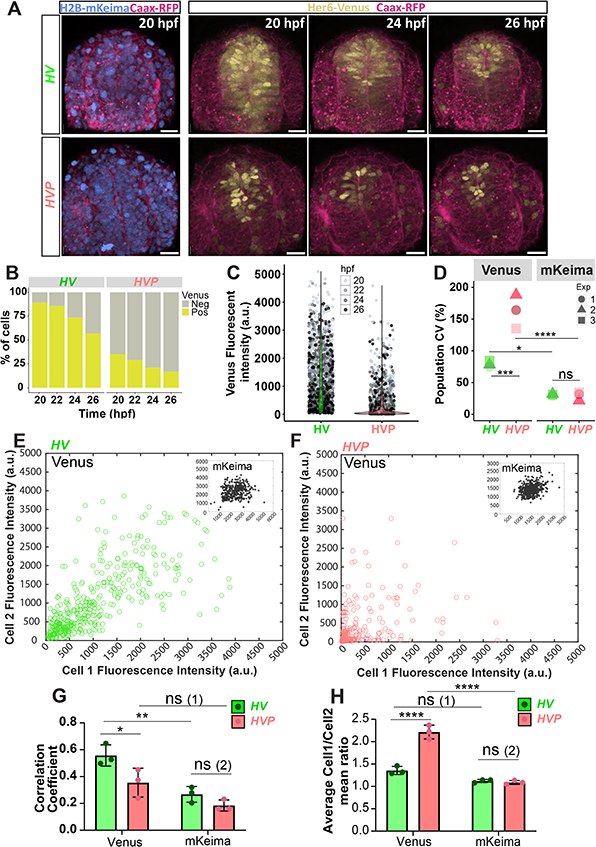
Spatial heterogeneity in HerG:Venus expression is altered in embryos with destabilised protein. **(A)** Representative examples of 3D reconstructed confocal images of the transversal view of the telencephalon from *Her6:Venus* (HV-top panels) and *Her6:Venus-PEST (HVP-bottom* panels) embryos between 20-26hpf showing HV and HVP expression (yellow), nuclei visualised with H2B-mKeima (blue) and membranes visualised with CAAX-RFP (red) (scale bar 30μm). **(B)** Percentage of HV and HVP positive cells over time from a single experiment shown in A (experiment 2, See Materials and Methods). Additional replicates quantified in **Figure S5A**. **(C)** Comparison of nuclear HV and HVP intensity expression from experiment 2 (dots indicate individual nuclei split by timepoint, violin plots indicate pooled distribution of nuclear intensities showing strong enrichment of low intensity levels in HVP (temporal breakdown for all 3 repeats is shown in **Figure S5C))**. **(D)** Comparison of population CV obtained from Venus (HV and HVP respectively) and H2B-mKeima expression from *HV* and *HVP* embryos observed in 3 replicates (marker indicate mean CV of all cells in the first 4 timepoints per embryo (on average 1630 and 1576 cells per *HV* and *HVP* embryo, respectively), statistical tests indicate two-way ANOVA with Tuckey’s multiple comparisons (temporal breakdown for all 3 repeats is shown in **Figure S5D**)). **(E-F)** Local intensity mapping of Venus (HV and HVP expression, respectively) in neighbouring cells observed in *HV* **(E)** and *HVP* **(F)** embryonic telencephalon imaged simultaneously at 24 hpf (dots indicate intensities in individual nuclei (Cell 1) that were paired with their nearest neighbour (Cell 2) based on 3D Euclidean distance (Materials and Methods), inset panels show H2B-mKeima intensity mapping from HVand *HVP* embryos, respectively). **(G)** Quantification of Pearson correlation coefficient computed from HV local intensity mappings (see **(E)** and **(F))** and corresponding H2B-mKeima (see **(E) and (F)** insets) observed between 20 to 26hpf in *HV* and *HVP* embryos imaged at the same time (bars indicate mean and SD of 3 biological repeats, dots indicate mean of 4 timepoints per embryo, statistical tests indicate 2-way ANOVA with Tuckey’s multiple comparison correction; ns (1) P=0.0936 and ns (2) P=0.5588). **(H)** Average intensity ratios observed in neighbouring cells showing Venus (HV and HVP, respectively) and H2B-mKeima quantified in *HV* versus *HVP* embryos between 20 to 26hpf. Neighbouring cells are paired based on minimal 3D Euclidean distance (see Materials and Methods) (bars indicate mean and SD of triplicate biological repeats, dots indicate mean of 4 timepoints per embryo, statistical tests indicate 2-way ANOVA with Tuckey’s multiple comparison correction and, ns (1) P=0.0901 and ns (2) P=0.9989). Statistical significance (ns): P >0.05, (*): P ≤ 0.05, (**): P ≤ 0.01, (***): P ≤ 0.001, (****): P ≤ 0.0001.

We then wanted to know if the reduced number of detected Venus(+) in *HVP* was due to global reduction of Her6 levels. To address this question, we combined cells from different time-points and from each experimental condition and looked at Venus levels at a population level. Overall, a large number of HVP cells had very low expression, as expected from the protein destabilisation, however unexpectedly some cells expressed Her6 at levels as high as HV **(Fig. 4C**-single experiment and **Fig. S5C-** all repeats and temporal breakdown). Further analysis of CV at population level confirmed that *HVP* has higher variability when compared to *HV* **(Fig. 4D**, and **Fig. S5D-** temporal breakdown). As expected, we observed minimal population heterogeneity in the nuclear control H2B-mKeima, which was significantly less compared to Venus and unchanged between *HV* and *HVP* embryos **(Fig. 4D)**. To ensure this effect was not due to the disproportionate presence of Venus(−) cells in *HVP* embryos, we also repeated this analysis in Venus(+) cells alone **(Fig. S5E-H)**, showing that Her6 protein expression was still more heterogenous in *HVP* **(Fig. S5H)**.

Overall, these results suggest that a decrease in Her6 stability increases both the temporal protein expression heterogeneity at single-cell level and the spatial protein expression heterogeneity at population level.

### Differences in Her6 expression between neighbouring cells increases in *HVP* embryos

To further examine the implications of the increased heterogeneity in Her6 level in *HVP* embryos, we looked at the relationship of Her6 expression between neighbouring cells in *HV* and *HVP* embryos at 24hpf **(Fig. 4E-H)**. For each cell, the closest neighbour was identified, and the intensity levels of Venus were plotted as paired sets with Venus intensity of a selected cell (cell 1) on the x-axis and its neighbouring cell (cell 2) on the y-axis **(Fig. 4E-F**, see Materials and Methods). In *HV*, cell 1 and cell 2 intensities produced an average correlation of approximately 0.56 **(Fig. 4E,G)** while *HVP* cells diverged and showed a lower average correlation of approximately 0.35 **(Fig. 4F,G)**. Thus, correlation in neighbouring intensities was significantly reduced in *HVP* compared to *HV* **(Fig. 4G)**. We also investigated differences in correlation coefficients at specific timepoints between 20 to 30hpf where we observed the same tendency for *HVP* to be less correlated than *HV* **(Fig. S6A,B)**. As expected, the nuclear control H2B-mKeima expression appeared more randomly distributed and similar between *HV* and *HVP* **(Fig. 4E,F** insets; **Fig. 4G** and **Fig. S6A** -timepoint breakdown). We further validated this result by an additional ratiometric analysis which corrects for these sources of technical variability (Materials and Methods). We calculated local intensity ratios between the paired data shown in **Fig. 4E,F** and reported averages per embryo at all times **(Fig. 4H)** and at specific stages **(Fig. S6C-E)**. Venus mean intensity ratios in neighbouring cells from *HV* were approximately 1.4 and significantly increased in *HVP* to >2 **(Fig. 4H** and **Fig. S6C-E)**. As expected, mKeima local intensity ratios appeared around 1 with no significant difference between *HV* and *HVP* **(Fig. 4H**, mKeima). In summary, neighbouring cells in *HV* have more similarity in local intensity resulting in increased positive correlation in terms of Her6 expression. Whereas in *HVP*, Her6 expression in neighbouring cells is less correlated and there is more than a 2-fold difference in Her6 expression in neighbouring cells.

The quantification of local and global differences in *HV* suggested that as protein stability is reduced, the heterogeneity in protein expression becomes increased and the relationship between neighbouring cells is altered. We investigated both single-cell autonomous and cell coupling mechanisms that could explain this effect using mathematical models.

### The unexpected increase in Her6 expression heterogeneity can be explained by cell-cell coupling

The destabilisation of the Her6 protein in *HVP* is found to increase cell-cell concentration differences when compared with *HV*. To understand how this increased heterogeneity might come about we generated three plausible networks of Her6 protein interaction and determined which models can lead to increased cell-cell differences when subjected to an increased protein degradation rate.

The simplest model, *Model 1*, consists of protein produced and degraded at a fixed rate within a single uncoupled cell, where Her6 expression does not affect expression in neighbouring cells **(Fig. 5Ai**, Materials and Methods). Through analysis of this model’s steady state concentration, we show that a population of cells would always decrease cell-cell concentration differences when the degradation rate is increased **(Fig. S7A)**. Furthermore, if we take the HV data and apply an increase to protein degradation according to this model (Materials and Methods), we do not reproduce the HVP distribution of intensities in the population **(Fig. 5Aii)**.

*Model 2* extends *Model 1* to include autoinhibition of the protein production rate, implemented with a Hill function **(Fig. 5B)**. One important parameter of the Hill function is the Hill coefficient, which when equal to one, gives classic Michaelis-Menten kinetics whereas high values give more non-linear, switch-like, kinetics. Given what is known about Her6 autoinhibition and the fact that it forms dimers, an upper limit of around 2 would be a realistic expectation for the Hill coefficient as it is linked to protein cooperativity (Weiss, 1997). As in *Model 1* we asked whether increased degradation could lead to increased cell-cell concentration differences, and we explored this over a range of Hill coefficient values. For Hill coefficient value less than 8, increased degradation always resulted in reduced cell-cell concentration differences **(Fig. S7B)**, whereas for values above 8, certain regions of parameter space showed an ability to cause increases in cell-cell concentration difference **(Fig. S7C)**. Given that a Hill coefficient of 8 is unrealistic, we determined that *Model 2* was also unlikely to account for the *HVP* phenotype.

**Figure 5:**
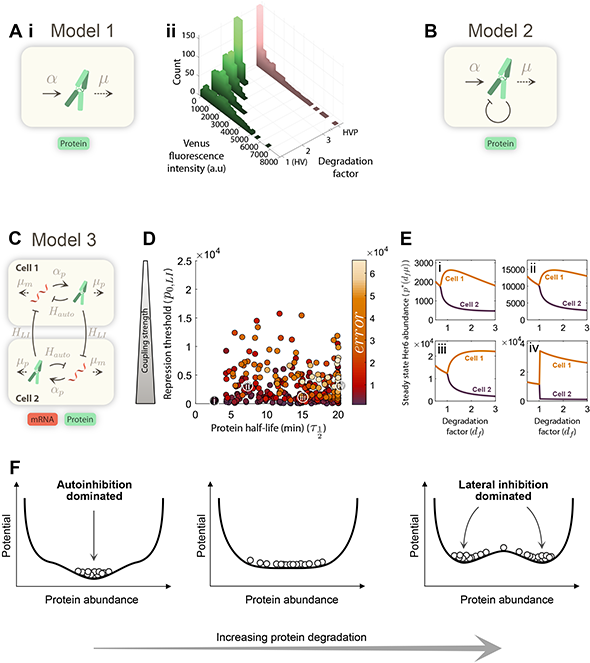
Coupled two-cell model of HerG protein in cells with normal and altered protein stability. **(A)** (i) Interactions used in model 1 which included production of protein (solid arrow, with rate a) and degradation of protein (dashed arrow, with rateμ). **(ii)** Model 1 applied to HV data taken from the first timepoint and each cell’s abundance is multiplied by 1/degradation factor (green histogram plots). Pink plot shows the HVP data from the first timepoint as a comparison. **(B)** The interactions used in model 2, which is the same as model 1 but with the addition of an autoinhibition loop. **(C)** The interactions used in model 3. *α_p_* and *α_m_* are protein and mRNA synthesis rates respectively, and *μ_p_* and *μ_m_* are the degradation rates. *H_auto_* is the Hill function used for the direct repressive action of Her protein on its own mRNA production. *H_LI_* is the lateral inhibition Hill function that represents coupling between cells via Notch signalling. Ranges of values used for all parameters are given in **Table S1**. **(D)** The results of the optimiser (described in methods) when run using model 3, where the two parameters of protein half-life and repression threshold are plotted against each other and the error for each parameter set is coloured by its final optimiser error. The optimiser was run 6340 times and filtered, leaving 291 parameter sets which are plotted. **(E)** Four examples parameter sets that correspond to the circles marked on (D) which show the steady state concentrations of the two cells over a range of degradation factors. **(F)** Illustration of how the bifurcation would look on a potential landscape as degradation increases (white circles with black edges represent individual cells).

Taken together, explorations of *Models 1 and 2* suggested that a cell autonomous mechanism could not explain the HVP phenotype. This prompted us to consider a non-cell autonomous mechanism that relies on cell-cell coupling between neighbouring progenitors in the tissue. In the neural developing tissue, progenitors expressing HES/Her are coupled through co-repressive interactions via Notch signalling, explored with theory and experimentation in (Biga et al., 2021; Hawley et al., 2022). Therefore, in *Model 3* we consider a two-cell model where single-cell dynamics are coupled together via lateral inhibition, representative of Notch-Delta signalling **(Fig. 5C)**. At the single-cell level, protein and mRNA levels are modelled as in (Monk, 2003) and cell-coupling is modelling as in (Biga et al., 2021). Lateral inhibition is capable of bifurcating expression between contacting cells where one cell adopts a low expression steady state and the other a high expression. The bifurcation occurs when the coupling strength between the two cells passes a certain threshold but here, we ask whether a similar type of bifurcation can occur not through coupling strength changes but when the rate of protein degradation changes instead.

We used a pattern search optimiser approach to efficiently explore parameter space and identify any regions with the desired phenotype in *Model 3*. This first required a range of reasonable parameter values to be defined over which the optimiser could run many times starting from random initial parameter values (Materials and Methods and **Table S1)**. Each parameter set tested by the optimiser was simulated twice, but with the second run of the model using an increased degradation rate of 1.1 x *μ*, where *μ* is the protein degradation rate. The 10% change in degradation rate roughly reflects the 12% observed change in the cycloheximide half-life experiments performed in MCF7 cells (where HVP has a faster degradation rate compared to HV) **(Fig. S4D**, Materials and Methods). We then measured how much the cell-cell concentration difference changes by defining the optimiser *error* function as

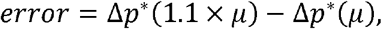

where Δ*p\** is the difference in the steady state concentration between the two cells.

With the optimiser set to move towards positive *error* values, parameter sets that satisfy the *HVP* phenotype of increases in cell-cell concentration difference were successfully identified (distributions for all parameters in **Fig. S7D)**. The identified parameter sets were found to increase cell difference through a bifurcation behaviour when a certain degradation rate is attained, as can be seen in **Fig. 5 Ei-iv**.

The analysis of the three models indicates that single-cell protein dynamics are unlikely to account for the observed distribution change in the *HVP* vs *HV* phenotype, but that Notch signalling, which couples the cellular dynamics, could account for the increase in heterogeneity when the system is subjected to increased protein degradation (summarised in **Fig. 5F; see discussion**).

### Embryos with altered Her6 dynamics and increased Her6 population heterogeneity are phenotypically normal at an early stage

So far, we have shown that destabilised Her6 leads to altered single-cell dynamics, accompanied by an increase in population heterogeneity due to cell coupling. Next, we sought to determine the phenotypic effects of these changes during the development of the telencephalon. We analysed the expression domains of *her6, ascl1, ngn1* that label GABAergic and glutamatergic progenitor cells in the sub-pallium and pallium respectively, as well as the post mitotic neurogenic marker *elavl3*, in *HV* and *HVP* at 24hpf (the mid-point of our live imaging). The spatial expression of *elavl3, ascl1* and *ngn1* was not altered in relation to *her6* in *HVP* embryos **(Fig. 6A-C** and **Fig. S8A)** but the volume of the *her6* domain was significantly reduced in *HVP* compared to *HV* **(Fig. 6D)**. This was not the case for *elavl3, ascl1 and ngn1*, where the volume of their expression domains was not significantly different, although we observed a tendency for those to be smaller in *HVP* **(Fig. 6D)**. These findings suggest that there are no marked changes in the domains of differentiation, as result of destabilising Her6, at this developmental stage.

**Figure 6.**
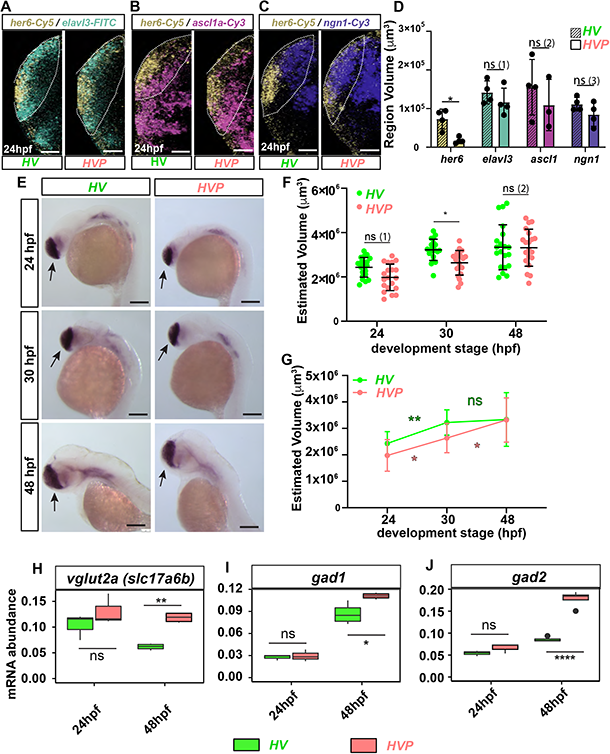
The phenotypic effect of destabilising Her6 protein is negligible in 24hpf embryos but becomes more evident by 48hpf. **(A-C)** Maximum intensity projection images of double WM FISH showing *her6/elav/3* **(A)**, *her6/asc/1* **(B)** and *her6/ngn1* **(C)** mRNA expression in the telencephalon of 24hpf *HV* versus *HVP* embryos (manual annotation denotes the telencephalon boundary in lateral view, scale bars 40μm). **(D)** 3D volumetric comparison of the *her6, elavl3, ascl1* and *ngn1* mRNA expression domain in *HV* versus *HVP* embryos (obtained from data **A-C**, see Materials and Methods) (bars indicate mean and SD of 3-4 images embryos, dots indicate domain volume per embryo, statistical test is two-tailed non-parametric Mann-Whitney, ns (1) P=0.2000, ns (2) P=0.6286 and ns (3) P= 0.4857). **(E)** WM Chromogenic ISH showing *foxg1* mRNA expression in *HV* and *HVP* at 24, 30 and 48hpf. *foxg1* is expressed across the telencephalon with the exception of the roof plate (arrows indicate the telencephalon region, scale bars 150μm). **(F)** 3D volumetric comparison of telencephalon size (calculated using length, depth and width measurements from data in **(E)**, see also **Fig. S8D)** in HVversus *HVP* from 24 to 48hpf (dots indicate individual embryos (n=20 per condition), bars indicate mean and SD from 20 embryos, statistical test is 2-way ANOVA with Sidak multiple comparison correction, lines across strain and time indicate significant contribution of strain (*HV* and *HVP)* and time to the variation in telencephalon size, ns (1) P=0.1104, ns(2) P>0.99). **(G)** Comparison of the changes in telencephalic volume over time (data from **(F))** (spots and bars indicate the mean and SD of telencephalon size between 20 embryos, statistical test is 2-way ANOVA with Sidak multiple comparison, ns:P>0.99). **(H-J)** Comparing mRNA expression of Glutamatergic marker *vglut2a*, GABAergic marker *gad1* and GABAergic marker *gad2* between *HV* (green) and *HVP* (pink) in dissected telencephalon of 24 and 48hpf embryos using qPCR (boxes indicate median and interquartile range from 3 or 4 biological repeats at 24 and 48hpf respectively with telencephalon from 20 embryos in each repeat, statistical test is 2-way ANOVA with Tuckey’s multiple comparison correctio n. **(H)** (ns) P=0.4729. **(I)** (ns): P>0.9999. **(J)** (ns): P=0.8174). Statistical significance (ns): P >0.05, (*): P ≤ 0.05, (**): P ≤ 0.01, (****): P ≤ 0.0001.

Given the tendency for smaller expression domains of the examined genes **(Fig. 6D)**, we suspected that there may be a more global difference in the size of the whole telencephalon. To measure the telencephalon we used chromogenic whole mount in situ hybridisation (WM ISH) against *foxg1* which is expressed in the entire telencephalon except for the roof plate (Materials and Methods). This was done using 24, 30 and 48hpf embryos to gain an insight into potential early and late phenotypes **(Fig. 6E** and **Fig. S8B-F)**. We measured depth, length and width from lateral and transversal images **(Fig. S8C-F)** and compared these indicators across multiple embryos in *HV* and *HVP*. There were no notable differences in depth and length at any stage **(Fig. S8D,E)** however, the telencephalon appeared wider in *HV* compared to *HVP* at 30hpf **(Fig. S8F)**. We also estimated changes in the overall volume of telencephalon **(Fig. 6F-G**, Materials and Methods) and identified a minor but significant reduction in the volume of *HVP* compared to *HV* at 30hpf **(Fig. 6F**, 30hpf). Despite small differences in telencephalon size at 30hpf, by 48hpf the telencephalon had regained its size and appeared normal in *HVP* when compared to *HV* **(Fig. 6F,G**, 48hfp). The recovery in size is likely due to differences in the telencephalic growth. Indeed, after an initial phase of rapid growth, the *HV* telencephalon does not increase between 30hpf and 48hpf, while the growth in *HVP* is gradual and steady between 24hpf and 48hpf **(Fig. 6G)**.

In short, even though there were clear differences in single-cell and tissue-level dynamics of Her6 expression in the telencephalon, there were only transient anatomical differences and no marked differences in subdomains. We next examined the expression of terminal neural differentiation markers.

### Changes in differentiation are observed in later stages of telencephalic development

We looked at the presence of terminally differentiated neurons, which are a clear readout of developmental output in the developing nervous system **(Fig. 6H-J)**. We compared the expression of three terminal neuronal markers between 24 and 48hpf in the *HV* and *HVP* telencephalon. We observed that the expression of *vglut2a* (Glutamatergic neuronal marker), *gad1* and *gad2* (GABAergic neuronal markers) were not different between *HV* and *HVP* at 24hpf but it was significantly higher in *HVP* at 48hpf **(Fig. 6H-J)**.

Taken together, our observations suggest that when Her6 is destabilised in single cells there is a compensatory mechanism at the population level, possibly mediated by the cell-cell coupling, that rescues Her6 protein level in some cells. In turn, this results in normal neural differentiation and telencephalic size, in spite of the premature loss of Her6 in most progenitor cells. However, such compensation is no longer effective at later stages of development, hence an increase in differentiation is observed.

## Discussion

In this manuscript, we have examined the role of protein biochemical properties in shaping Her6 dynamics at single-cell level, and the consequences of altering such properties at the tissue level, focusing on the development of the zebrafish telencephalon. We have discovered that reducing Her6 protein stability, by an in-frame fusion of a destabilising PEST domain to the endogenous protein, reduces the Her6 protein levels overall and causes for the Her6 expressing domain in the telencephalon to shrink prematurely over the course of development. However, the phenotypic effect was mild and only evident at late stages in development, suggesting that a compensatory mechanism may be operating to “rescue” normal development.

HES/Her proteins tend to be unstable and protein instability is a prerequisite for oscillations to occur (Hirata et al., 2002) and indeed, Her6 is naturally an unstable protein. The addition of a PEST domain had a small effect in the overall stability of the protein, which was revealed when the protein was introduced in a heterologous system (a mammalian cell line); yet, the effect on the protein dynamics was pronounced. This demonstrates that the dynamic output of the HES/Her negative feedback loop is carefully balanced around the stability values of the protein. Thus, manipulations of protein stability can join previous manipulations of biological time delays (Takashima et al., 2011; Harima et al. 2013; Ochi et al., 2020; Shimojo et al., 2016) and mRNA stability/translation (Bonev et al., 2012; Ochi et al., 2020; Soto et al., 2020; Tan et al., 2012) as an effective way to change the output of a regulatory negative feedback loop, as theoretically predicted (Lewis, 2003; Monk, 2003).

Alongside the expected reduction of protein abundance in most neural progenitor cells, due to reduced protein stability, we have made two counter-intuitive findings; first, we observed an increase in the proportion of cells with oscillatory Her6 and furthermore, these had an increased amplitude than normal, indicating more and better-quality oscillations. Second, some progenitor cells appeared to maintain Her6 expression at the wild-type level in *HVP* embryos. As the work was carried out in homozygous stable lines of fish, we can exclude the possibility of genetic mosaicism as the cause for these findings. Thus, these unexpected results consequently meant that there was an increased heterogeneity of Her6 protein levels at the population level and increased differences between neighbouring cells. The latter seen as increased CV and decrease in correlation between neighbouring cells in snapshot measurements.

Her6, like other HES/Her genes, is expressed in proliferating progenitor cells and in the mouse, combinations of HES gene knockout leads to failure of progenitor maintenance and premature differentiation (Ochi et al., 2020). Thus, we expected that the premature depletion of Her6 expressing cells in *HVP* embryos will result in premature differentiation. Surprisingly, the telencephalon appeared normal, at least with the criteria employed here. That is early differentiation markers (*elavl3, ngn1* and *ascl1)* and overall size of the telencephalon (measured based on *foxg1 expression)* at an early stage of development (24hpf). At later stages (48hpf), there was an increase in some terminal differentiation markers. The late manifestation of a (mild) phenotype contrasts with the early onset of Her6 expression, which has been detected as early as 11hpf (see **Fig S1)**.

We surmise that cells that maintain normal Her6 levels somehow compensate for the premature loss of Her6 in other cells, perhaps by undergoing extra proliferation at the expense of differentiation. This is supported by the growth curve of the *HVP* telencephalon which appears to “catch up” with the HV telencephalon between 30 and 48hpf. It is tempting to speculate that the increased proportion of high amplitude oscillations is also part of a compensatory mechanism. The increase in oscillatory activity is consistent with our previous prediction that increased protein degradation rate would allow cells to enter an oscillatory regime (Manning et al., 2019).

How do high Her6 expressing cells arise if the protein is globally destabilised? The answer to this question is not intuitive, therefore we employed mathematical modelling to get predictive insights into the potential mechanisms. Our modelling showed that destabilisation of Her6 affects the regulatory network in such way that a bifurcation occurs between neighbouring cells leading to population-level increase in low/off cells, while some cells retain normal levels of Her6 expression. This effect is not observed when Her6 is modelled in single cells; in other words, the bifurcation of the population into high and low Her6 expressing cells in *HVP* embryos arises as an emergent property of cell-cell coupling. A prerequisite is that such coupling mechanism should have the ability to affect Her6 expression in neighbouring cells and this condition is easily satisfied by lateral inhibition via Notch signalling which is known to generally regulate expression of HES/Her genes.

However, the dependence on Notch signalling, the precise nature of the response (activation or repression) and the precise type of Notch receptor/ligand involved, are variable among Hairy/E(spl) family members (e.g. Hans et al., 2004; Schmidt et al., 2013; Tseng et al., 2021). Therefore, experimental validation of the dependence on Notch signalling will be needed in future studies. Additonally, other signalling pathways that are present in the forebrain, such as Shh (Wilson and Rubenstein, 2000; Danesin et al., 2009) may contribute to the regulation of Her6, as is the case for Hes1 (Wall et al., 2009)

The underlying cause of the bifurcation is not intuitive and appears to depend upon an interplay between both Her6 autoinhibition and lateral inhibition. Indeed, our 2-cell model (Model 3) fails to recapitulate the *HVP* phenotype when it is only run in the presence of lateral inhibition but without autoinhibition. Mechanistically, we propose that in the system, initially the autoinhibition has a stronger influence than lateral inhibition and maintains both cells at a single high expression value **(Fig. 5F)**. Then, bifurcation arises as degradation increases and protein abundance drops, leading to lateral inhibition dominating **(Fig. 5F)**. For the cells to be primed to bifurcate and switch from a single autoinhibition attractor state to the two attractor states of lateral inhibition, there is a minimum level of coupling strength required, which is reflected by the fact the optimiser finds solutions within a restricted range of coupling strengths in **Fig. 5D**. Although this proposed model is an attractive possibility, presently, we cannot exclude the possibility that phenotypic compensation occurs wholly or partially by a different mechanism such as, for example the ectopic expression of one of the other Her family genes, as it has been described in the mouse (Ochi et al., 2020).

In conclusion, altering Her6 protein stability has a knock-on effect on altering single-cell dynamics but cell-cell coupling may act to rescue developmental abnormalities. Such rescue mechanism is not perfect, because at a later stage increased differentiation is observed, suggesting that the limits of compensation have been reached. We suggest that emergent properties of coupling single cell Her6 oscillations not only control the spatiotemporal order of neurogenesis, as we have shown before (Hawley et al., 2022), but also provide a mechanism for phenotypic robustness to the whole tissue when single-cell dynamics are altered.

## Materials and methods

### Research animals

All animal work was done in line with the conditions of the Animal (Scientific Procedures) Act 1986 and under the UK Home Office project licence (PFDA14F2D). Animal handling was carried out by personal licence holders.

### Characterisation of the knock-ins by step-wise Polymerase chain reaction (PCR) and sequencing

Genomic DNA was extracted from 2-4 Days post fertilisation (dpf) embryos using NP40/Proteinase K (PK) extraction method. 20μl of extraction solution (25μl of PK (NEB) /ml of NP40 lysis buffer) was added to each embryo and incubated at 55°C for 3-4 hours, followed by enzyme deactivation at 95°C for 15 minutes.

All PCR reactions for characterising were done using the Phusion High-Fidelity DNA Polymerase kit (Thermo Fisher) based on manufacturer instructions using the primers in **Table S2**. For sequencing, in some cases amplicons were cloned using Zero Blunt TOPO PCR cloning Kit (Invitrogen) using manufacturer protocol. Positive clones were sequenced using M13 primers included in the kit. In other cases amplicons were directly sequenced using the same primers used for amplification.

### Molecular cloning

To generate PCS2-Her6-Venus-HA and PCS2-Her6-Venus-PEST-HA plasmids, Venus sequence was amplified from Her6-Venus (HV) CRISPR donor designed by Soto et al. (2020) using the following primers with inclusion of Eco-RI restriction sites **(Table S3)**.

The purified Venus-EcoRI amplicon was cloned into pCR-Blunt 11-TOPO (Invitrogen) using manufacturer guidelines. Plasmid was purified from selected colony using the QIAprep Spin Miniprep Kit (Qiagen), checked for insertion using restriction enzyme digest with Eco-RI (NEB) and sequenced (Eurofins, LIGHTRUN).

To generate both, pCS2-her6-venus-HA and pCS2-her6-PEST-venus-HA vector, we excised out PA GFP from PCS2-Her6-PA GFPHA and PCS2-Her6-PA GFP-PEST-HA with Eco-RI digestion. The digested plasmid was resolved using agarose gel electrophoresis and purified from the gel with QIAquick Gel Extraction Kit. Vectors were treated with Antarctic Phosphatase (NEB) to prevent re-ligation. PCS2-Her6-HA and PCS2-Her6-PEST-HA (vectors) and Venus-EcoRI (insert) were assembled using T4 DNA ligase (NEB) using the manufacturer protocol. The resulting colonies were examined by colony PCR using MyTaq Red mix (Meridian Bioscience - BIO-25043) with Venus primers shown in **Table S4**. Positive clones for pCS2-her6-venus-HA and pCS2-her6-PEST-venus-HA were sequenced and purified using PureLin HiPure kit (ThermoFisher).

### Chromogenic and Fluorescent Whole mount In situ hybridisation

*foxg1* antisense probes labelled with Digoxigenin (Dig) were generated using foxg1-pBluescriptII construct kindly gifted by Corine Houart and chromogenic Whole-mount In-situ hybridisation (WM ISH) against *foxg1* was carried out as previously described by Thisse (Thisse and Thisse, 2008) The antibody used to detect the riboprobes was AP-anti-DIG (Roche Cat.N: 11093274910, 1:500). For Fluorescent WM ISH (WM FISH), the her6-Dig, her6-Dinitrophenol (DNP), Elavl3-Fluorescein (FITC), *venus-FITC* were generated from constructs previously described in (Soto et al., 2020), while *ngn1*-Dig and *ascl1-Dig* probes were generated from constructs kindly gifted from Dr Laure Bally-Cuif. Tyramide amplification was used after the addition of probes and horseradish peroxidase conjugated antibodies as described by (Lea et al., 2012), Anti-DIG-POD (Roche Cat. N: 11207733910, 1:500) and Anti-FITC-POD (Roche Cat. N. 11426346910, 1:500).

WM ISH embryos were submerged in Glycerol on glass dishes, positioned to desired orientation and imaged using a Leica M165 FC Stereo microscope. All image processing and measurements from ISH were done using FIJI measurement tool.

WM FISH embryos were imaged using the Upright LSM 880 Airyscan microscope in Fast Airyscan mode with a W Plan-Apochromat 20x/1.0 DIC (UV) VISIR M27 75mm lens. The first step of processing was done using default Airyscan Processing setup in ZEN Black. Further analysis of domain sizes was done by generating a mask of the signal using Surfaces tool of Imaris 9.3.1 software. The surface grain size was between 2-3μm and the threshold was set manually for each gene in each embryo based on the best representation of the signal. The volume of the mask used for the analysis was automatically quantified by the Imaris software.

### Whole mount immunofluorescence

Whole mount IF was adapted from (Mendieta-Serrano et al., 2013). Embryos were collected and fixed with 4% Formaldehyde-PBS. They were washed 2×10 mins in blocking solution (PBS, Bovine serum albumin (BSA) 0.1% (Sigma Aldrich - A7906) and Triton X-100 (TX-100) 1%) and 2×10 mins in PBS-TX while rocking at RT. For permeabalisation, embryos were treated with Proteinase K (10μg/ml in PBS) for 10 mins for 24 hours post fertilisation (hpf) embryos or 13.5 mins for 48hpf embryos and then washed 2×5 mins in PBS-TX. Embryos were blocked for 2 hours at RT with rocking. After blocking, embryos were incubated with chicken polyclonal anti-GFP antibody (abcam - ab13970, 1:300) and rabbit polyclonal anti-Phospho-Histone H3 (Ph3) antibody (Sigma Aldrich 06-570, 1:500) in blocking solution overnight (ON) at 4°C. Primary antibodies were washed off the following day 1×20 min in blocking solution and further 5×10 min washes at RT with rocking. Following from this, they were incubated with Alexa Fluor 568 goat anti-rabbit antibody (Thermo Fisher - A11011, 1:500) and Alexa Fluor 488 donkey anti-chicken antibody (Jackson lmmunoresearch - 703-545-155, 1:500) in blocking solution for 3 hours at RT while rocking. They were washed for 15 and then 10 mins in blocking solutions and a further 10 mins in PBS-TX. For nuclear staining, embryos were incubated in DAPI (Thermo Fisher - 62248) at a final concentration of 5μg/ml at 4°C ON while rocking. On the following day, DAPI solution was removed and embryos were washed 3×10 mins with PBS-TX.

### Quantification of volume from WM ISH datasets

For each condition, 20 embryos were imaged in both transversal and lateral orientations. Using the FIJI measure tool, the length (L) and depth (D) of the telencephalon was estimated from the lateral images and the width from the transversal images **(Fig. S8D)**. The values were converted from arbitrary units to μm using the scale bar recorder on the images. The volume of the *foxg1* expression domain (calculated by multiplying the three dimensions DxLxW) was used as a proxy of the telencephalic volume.

### Live imaging

mRNA for cellular markers was generated in vitro using mMESSAGE mMACHINE™ SP6 Transcription Kit (Thermo Fisher) and purified with MEGAclear Transcription Clean-Up Kit (Thermo Fisher). All embryos were injected with approximately 1nl of solution consisted of mRFP-Caax (40ng/μl), H2B-mKeima (40ng/μl) and 0.05% Phenol red as described by (Soto et al., 2020).

Embryos were mounted 1 hour prior to imaging in a 50mm glass bottom dish (MatTek Corporation), positioned face-up for a transverse view in 1% LM agarose with MS222 (final concentration 160ng/ml). After 1 hour of setting at Room Temperature (RT), the dish was filled with embryo water supplemented with MS222 (final concentration 160ng/ml) and N-Phenylthiourea (PTU) (0.045% stock-Sigma Aldrich) which was circulated with a peristalsis pump (Harvard Apparatus). All live imaging was done at 28°C on a Zeiss Upright LSM880 Airyscan microscope in Fast Airyscan mode. We used W Plan-Apochromat 20x/1.0 DIC (UV) VISIR M27 75mm lens, 3x zoom, Image size 139×139μm, Z: 81-86 and Z size: 38.4-43.4 μm. The three channels were imaged sequentially and the filters used were Channel 1:BP 420-480 + LP 605, Track2: BP420-480 + BP495-550 and Track 3:BP 420-480 + LP 605. The lasers used were Channel 1: 561nm (between 2-10% for mRFP), Channel 2: 458nm (between 6-10% for mKeima) and Channel 3: 514nm (between 6-13% for Venus). Images were captured every 6 minutes for 6-12 hours. The Airyscan files were processed using the automatic Airyscan processing tool of the ZEN black software.

### Nuclear segmentation and single cell tracking from live imaging data

We used the Spots tool on Imaris 9.3.1 to automatically identify cells in the domain of interest based on the nuclear marker H2B-mKeima. The automated detection was checked manually for Venus or H2B-mKeima expressing nuclei that may be missed due to the signal quality.

For the population analysis, both Her6:Venus and H2B-mKeima mean fluorescence intensities were extracted from these cells from selected timepoints, starting from the first frame of imaging at 20 hours post fertilisation (hpf) and continuing at 2-hour intervals up to 26hpf.

For distinguishing between Venus (+) and Venus (−) a threshold was determined for each embryo. For this, we first calculated the median value of background intensity in each individual embryo as a starting threshold. Then we manually examined cells with expression above and below this initial threshold and adjusted it if needed to represent the boundary between visible Venus(+) and Venus (−) cells.

For single cell traces analysis, single nuclei were tracked and analysed using methods described previously by (Soto et al., 2020) using Imaris 9.3.1. In short, we first subtracted the H2B-mKeima signal (Ch2, nuclear marker) from the Caax-mRFP signal (Ch1, membrane marker) with the “arithmetic tool” to remove background signal within the boundaries of each cell. The resulting channel (Ch4) was then subtracted from the Her6:Venus signal (Ch3) to segment as distinguishable nuclei. We then used the ‘Spots’ (5μm in XY and Z diameter) and ’Track over time’ functions to curate tracks of individual cells over time using a combination of automatic and manual tracking. In cases where automatic tracking was used, all final tracks were manually checked to ensure the same cell is tracked over time and there has been no mixing between neighbouring tracks. We also generated background tracks to measure background fluorescence. All statistics were exported from Imaris and processed. All track data were exported from Imaris 9.3.1. Since the Imaris 9.3.1 version does not distinguish between daughter cells connected to the same mother cell when exporting the intensity values, we used the Track Reconstruction (tRecs) Python script (https://github.com/TMinchington/tRecs) to connect the tracks of daughter cells and re-form the cell families in the exported data. Tracks shorter than 3h were rejected and for cells that divided only one track was considered (out of the two daughter cells).

### Detection of oscillatory and aperiodic fluctuating activity from time-series

Dynamic analysis of single cell tracks was performed using the method developed by (Phillips et al., 2017) and adapted for zebrafish single cell tracks by (Soto et al., 2020). In summary, we first normalised Her6:Venus to the H2B-mKeima signal by division **(Fig. 2Giii; HV/H2B Normalised)** to correct for any fluctuations in Her6:Venus that were caused by global changes in transcription and translation or technical issues during image acquisition. Next, since the analysis pipeline is most accurate in detecting oscillators in absence of long-term trends (Phillips et al., 2017), we detrended the Her6:Venus/H2B-mKeima from long-term trends of 4.5 hours **(Fig. 2Giv; HV/H2B normalised-detrended)**, a recommended value representing 3 times the expected period (Phillips et al., 2017) as estimated in Her6 hindbrain progenitors (Soto et al., 2020).

The detection method uses two competing Gaussian Process covariance models corresponding to (1)-random aperiodic fluctuating (OU) or (2)-noisy periodic wave (OUosc). During the analysis, the experimental time-series data is used first to infer optimal parameters for both models separately and the probability of the data under each model is computed. Time-series collected from background are used to calibrate technical noise. The confidence in the data being oscillatory is expressed as a Log likelihood ratio (LLR) of the probability of the data under the oscillatory model over the non-oscillatory model. To select the LLR threshold for classifying oscillators and controlling the False discovery rate (FDR), a set of synthetic data is generated from the OU model and synthetic aperiodic LLR scores are calculated.

By comparing the LLR score from the non-oscillatory synthetic data with LLR from the experimental data, we selected the LLR threshold suitable for our data with the stringent FDR of 3%. In order to ensure a fair statistical comparison, the datasets were paired in FDR as well as statistical testing. Specifically, in the comparison of HV/H2B versus H2B-mKeima **(Fig. 2)** and the comparison of HV to HVP **(Fig. 3)** the corresponding datasets were analysed as paired. This produced a similar % osc in HV **(Fig. 2J** and **Fig. 3F)**. The aperiodic technical control % H2B-mKeima passing at the same LLR threshold was approx. 5% across all pairings. While the imposed FDR is 3% in the synthetic data, a 5% passing rate of the technical control signal H2B is within accepted tolerance values.

For the amplitude of fluctuations and oscillations, peaks and troughs in the HV/H2B normalised tracks were identified using the Hilbert transform applied to detrended HV/H2B. Subsequent peaks and troughs were paired and their position was used to identify corresponding intensities in the raw HV/H2B signal and calculate fold-change (peak/trough) in the raw data (Manning et al., 2019). We report maximum peak to trough fold-change per track throughout.

### Intensity analysis from fixed point nuclear detection of neighbouring cells

The 3D coordinates of each detected nucleus in the cell population snapshot of selected timepoints were used to measure the inter-nuclear distance at approximately 2.3μm with no differences between HV (2.314 ±0.1326μm) and HVP (2.343 ±0.1388 μm). The closest neighbour to each nucleus was identified based on minimal 3D Euclidean distance and the intensity levels of Venus and mKeima were stored in paired datasets referred to as cell-cell intensity distributions. Some signal variability was associated with Z depth due to imaging artefacts or other uncharacterised expression gradients. To circumvent their effect on the analysis, ratiometric analysis was performed where the ratio between mean fluorescence of each selected nucleus was calculated in relation to its nearest neighbour in each embryo referred to as cell-cell intensity ratios.

### Telencephalon dissection and RNA extraction

To dissect the telencephalon, a petri dish was coated with 1% agarose in embryo water and once set, punctured with a 10μ1 pipette tip to generate small holes. 24 or 48hpf embryos were anaesthetised with MS222 and placed in the small holes generated on the agarose plate to prevent them from moving.

Embryos were visualised with intermediate contrast on a stereo microscope and the telencephalon was dissected using a Gastromaster microdissection machine at the “Highest” settings. The dissected tissue was transferred with a 10μl pipette tip coated with low-melting (LM) agarose (to prevent it from sticking to the plastic) into 500μl of ice-cold Trizol. 20 dissected telencephalons were pooled in each sample.

For RNA extraction, tissues were dissociated by pipetting. 100μl of chloroform was added to each sample and incubated at RT for 5 mins. Samples were centrifuged for 15 minutes at 21000g and supernatant was transferred to a fresh tube. 0.5μl of Glycoblue (Invitrogen) was added to facilitate RNA precipitation and visualisation along with 250μl of Isopropanol. Samples were incubated at - 20°C overnight (O/N). They were then centrifuged at 21000xg for 1 hour and supernatant was removed. Pellet was washed with 500μl of 70% EtOH and centrifuged for 5 mins at 21000g. Pellets were air dried for ~5 mins at RT and resuspended in 7-10 μl of RNAse free water (Invitrogen).

### Quantitative PCR

To remove any potential DNA contamination, RNA samples were first treated with RQ1 RNase-Free DNase (Promega) system according to manufacturer’s instructions. cDNA was generated using Superscript Ill Reverse Transcriptase kit (Invitrogen). For qPCR reaction, the probes shown in **table S5** were used with TaqMan Universal PCR Master Mix (Applied Biosystem) according to manufacturer’s instructions. 5ng of cDNA was used in a 5μ1 reaction and each sample was run with 2 technical repeats. qPCR was carried out using a 96-well StepOnePlus Real-Time PCR System with quantitation (comparative CT) and TaqMan experimental setup as a standard run.

CT values were analysed as follows: The technical repeats for each sample were checked to ensure there are no differences larger than 1 unit and were then averaged for each gene per sample. Actinb1 (a highly expressed gene and an established qPCR control) and Tmem50a (recommended housekeeping gene for comparing zebrafish developmental stages (Xu et al., 2016)) were used as controls. The CT values for both housekeeping genes in each sample were averaged to give a single housekeeping CT value. _Δ_CT was calculated by subtraction of housekeeping CT from the CT value of each gene of interest. ΔΔCT was calculated as 2^ΔCT^.

### Cyclohexamide chase experiments in MCF7 cells

Human breast cancer MCF7 cells (purchased from ATCC and regularly screened for mycoplasma contamination) were transfected with Lipofectamine 3000 (Invitrogen) according to manufacturer’s instructions. 25K-35K cells were plated in one quarter of a 4-quarter glass bottom dish (Greiner) in DMEM media (Sigma Aldrich) supplemented with 10% fetal bovine serum (Gibco). When cells reached 60-70% confluency (often after 24h) they were transfected with 500ng HV or HVP plasmid. 2 days post transfection cells were treated with 5μg cycloheximide (Sigma-Aldrich) and imaged straight after on Inverted LSM 880 Airy/FCS Multiphoton (NLO) microscope with a Plan-Apochromat 20x/0.8 M27 objective and image acquisition every 17 minutes using tile scanning, for about 22 hours. Two separate detectors were used for high and low Venus levels, since plasmid transfection results in highly variable levels of expression.

Maximum intensity projections of all images were generated using FIJI and data were randomised. Cell tracking was performed blindly on the randomised data on Imaris 9.3.1 using the Spots function with spot diameter of 11.9 μm. The mean Venus intensity from all spots was extracted and when signal was saturated with the detector capturing lower levels of Venus expression, the mean intensity from the other detector was used. Cells with saturated signal were excluded.

To estimate the protein half-life, the Venus expression tracks for HV or HVP were pooled together for each biological experiment (n=3) and GraphPad Prism 9.3.1 was used to fit one phase decay exponential curve to each data set.

### Statistical analysis

All data were processed using R-4.1.3. Main R packages used were readr 2.1.2, tibble 3.1.6, dplyr 1.0.8, tidyr 1.2.0, ggplot2 3.3.5 and tidyverse 1.3.1. All statistical analysis was done using GraphPad Prism Version 9.3.1. Detailed test analysis are included in figure legends.

### Quantification of heterogeneity by Coefficient of variation

Coefficient of variation (CV) is a metric that measures the relative variability around the mean in the data. This is a very useful tool for comparing variability in grouped data where different groups do not have comparable means like Venus and mKeima or HV and HVP in the context of the present work. CV is calculated by the division of standard deviation (SD) by the mean. Two types of CV were used:

1. Population CV (used in **Fig. 2B, Fig. S5D,F)** was obtained by calculating the mean intensity of all cells detected in each snapshot followed by calculating the SD of intensities of all cells in that population. Next, the SD was divided by the mean for each snapshot resulting in a single CV value for that snapshot. In cases where several timepoint snapshots are pooled, the mean, SD and CV were calculated for the whole pool (used in **Fig. 4D, Fig. S5H)**.
2. Time-series CV (used in **Fig. 2E, Fig. 3C** and **Fig. S4A)** was obtained by calculating the mean intensity of each single tracked cell over the duration of its time-series as well as the standard deviation of all timepoints. Next, the standard deviation was divided by the mean resulting in time-series CV.

### Mathematical modelling

#### Model 1: Uncoupled cells with no protein auto-regulation

Model 1 is described by

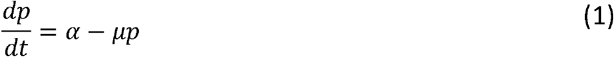

where *p* is protein abundance, *t* is time, *a* is the protein production rate, and *μ* is the protein degradation rate. The steady state can be analytically found by setting 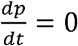, which yields

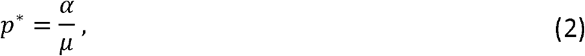

where *p\** is the steady state protein abundance. If we assume that a range of production and degradation values (*α* and *μ*) account for the heterogeneity in Her6 expression at tissue level, and that Eq. (2) describes how steady state will change with altered degradation, it can be shown that cell-cell concentration difference will always decrease with increased degradation, and therefore rule out model 1 as being able to account for the HVP phenotype.

If we take any two cells in the population, *i* and *j*, where *i* has a higher steady state expression than *j*

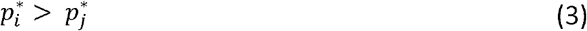

then for cell-cell concentration difference to decrease when *μ* is increased, the higher expressing cell *i* must have a more negative change in expression than cell *j*

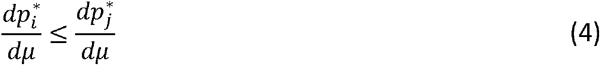

In the case of model 1, condition (4) holds for any two steady state concentrations, because the steady state decreases faster for high steady state values than low steady state values, as can be seen by the exponentially decreasing curve in **Fig. S7A**. The gradient of this line which is given by

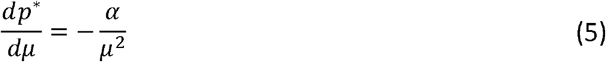

is a negative function that always increases with *μ*, **(Fig. S7A)**. This exponential decrease can be applied directly to the HV experimental data to show how the distribution would be expected to change under the assumption of model 1 by multiplying each experimental value by 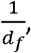 where *d_f_* is a degradation factor indicating the amount *μ* has increased, and this is shown in **Fig. 5A ii**.

#### Model 2: Uncoupled cells with auto-inhibition

Model 2 is the same as model 1, but includes an additional autoinhibition interaction in the form of a Hill function, which is more representative of known Her6 regulation and is described by the equation

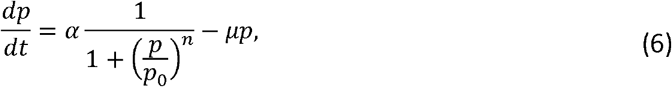

where *p_0_* is the repression threshold, which defines the protein abundance at which the protein production rate will be half maximum, and *n* is the Hill coefficient which define how switch like the self-repression will be.

If a value of *n* = l is used in Eq. 6, then the same type of analytical approach used for model 1 again finds that increasing μ always results in reduced cell-cell concentration difference. For *n* = 2, numerical analysis was used to explore the model and a range of *p_o_* versus 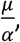 was explored and steady states plotted in **Fig. S7B i**. It can be seen that for each value of *p_0_* the steady state value has the same general shape of that in model 1, where steady state concentration decreases exponentially with increasing 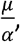 implying that again cell-cell concentration differences will always decrease with increased degradation in this model. This is confirmed by plotting the gradient 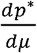 **(Fig. S7B ii)**, which is found to be a negative and always increasing, and therefore does not describe the phenotype.

At higher Hill coefficient values, starting at around *n* = 8, condition (4) does not always hold and the gradient of the steady state abundance **(Fig. S7C i)** is not always an increasing function as can be seen more clearly in **Fig. S7C ii**. At high Hill coefficients this implies that is possible for higher expressing cells to reduce expression at a slower rate than lower expressing cells in response to an increased degradation rate, which would lead to an increased cell-cell concentration difference across a population of cells. However, the value of the Hill coefficient being so high is extremely unlikely due to the high level of non-linearity produced by a relatively simple interaction (Weiss, 1997). Therefore, within reasonable bounds for the Hill coefficient, model 2 is unlikely to account for the HVP phenotype.

### Model 3: Coupled 2-cell model with auto-inhibition

Model three extends to a two-cell model coupled together via lateral inhibition, representative of Notch-Delta signalling and is adapted from (Biga et al., 2021) and also includes time-delays and mRNA dynamics.

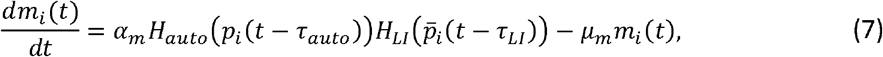

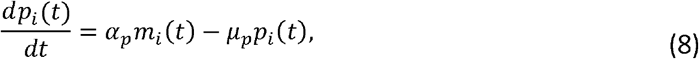

where *m_i_(t)* and *P_i_(t)* are the mRNA and protein abundance in cell *i* (as this is a two-cell model, *i* = 1, 2). *H_auto_* and *H_LI_* are the autoinhibition, and lateral inhibition Hill functions which in full are

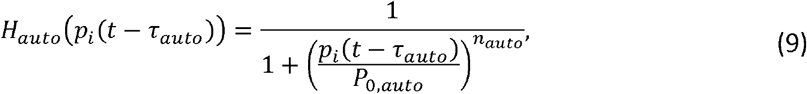

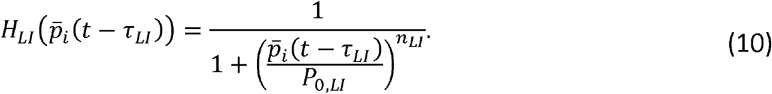

All parameters are defined in **Table S1**.

The optimiser approach used to identify increased cell-cell concentration differences in model 3 used MATLAB’s pattern search algorithm. The *error* function used was

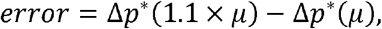

where Δ*p\** is the difference in concentration between the two cells, and the optimiser was set to maximise the *error*. The optimiser was run 6340 times from random initial parameters each time, and filtered for *error* > 1000 to results were causing significant increases in cell-cell concentration differences and *coherence* < 0.3 to remove oscillatory solutions from the results, which resulted in 291 accepted parameter sets (coherence measured in the same way as described in (Phillips et al., 2016). Histograms generated from the optimised parameter sets **(Fig. S7D)** indicate the most common values that are found to generate increases in cell-cell concentration differences.

## Data availability

The mathematical model (Materials and Methods-Mathematical modelling) was implemented in Matlab. It is deposited and available on GitHub at https://github.com/PapalopuluLab/Doostdar2023.

